# The structure of the Pro-domain of mouse proNGF in contact with the NGF domain

**DOI:** 10.1101/333070

**Authors:** Robert Yan, Havva Yalinca, Francesca Paoletti, Francesco Gobbo, Laura Marchetti, Antonija Kuzmanic, Doriano Lamba, Francesco Luigi Gervasio, Petr V. Konarev, Antonino Cattaneo, Annalisa Pastore

## Abstract

Nerve Growth Factor (NGF) is an important neurotrophic factor involved in the regulation of cell differentiation, maintenance, growth and survival of target neurons. Expressed as a proNGF precursor, NGF is then matured by furin-mediated protease cleavage. Increasing evidence suggests that NGF and proNGF have distinct cellular partners which account for different functional roles. While the structure of mature NGF is available, little is known about the structure of the pro-domain within the context of proNGF because the dynamical and structural features of the protein have so far prevented its structure determination. We have exploited a new hybrid strategy based on nuclear magnetic resonance and modelling validated by small angle X-ray scattering to gain novel insights on the pro-domain, both in isolation and in the context of proNGF. We show that the isolated pro-domain is intrinsically unstructured but has a clear tertiary structure propensity and forms transient tertiary intramolecular contacts. It is also able to interact, albeit weakly, with mature NGF and has per se the ability to induce growth cone collapse, indicating functional independence. Based on paramagnetic relaxation enhancement data and advanced molecular modelling, we have then reconstructed the overall properties of the pro-domain in the context of proNGF and showed that it has a compact structure. Our data represent an important step towards the structural and functional characterization of the properties of proNGF and its pro-domain.

## Introduction

Nerve Growth Factor (NGF) (Levi-Montalcini, 1987), the prototype member of the neurotrophin family of neurotrophic factors, is involved in the regulation of differentiation, maintenance, growth and survival of specific populations of peripheral and central neurons, as well as of non-neuronal cells (Huang and Reichardt, 2001). Mutations of NGF gene are linked to hereditary sensory autonomic neuropathy type 5 (HSAN5) (Capsoni, 2014), while alterations of NGF signalling have been associated to chronic and inflammatory pain (Pezet and McMahon, 2006) and neurodegeneration (Cattaneo and Calissano, 2012). It is secreted as a homodimeric precursor (proNGF) which is then proteolytically processed by furin to yield mature NGF (Seidah et al., 1996). Accumulating evidence shows that NGF and proNGF have distinct functions (Chao et al., 2006). The precursor is the more abundant form in central nervous system tissues, whereas mature NGF is barely detectable (Fahnestock et al., 2001). ProNGF can also be cleaved extracellularly as well as intracellularly, yielding the mature form and the pro-domain (Gibon and Barker, 2017). Therefore, the cleaved pro-domain exists *in vivo*, together with uncleaved proNGF and mature NGF, in a tightly regulated homeostatic equilibrium whose alteration can have profound consequences for neurodegeneration pathologies in the brain (Capsoni et al., 2011; Iulita and Cuello, 2014). It has also been reported that, in HEK TrkA stable cells, the pro-domain binds to TrkA at a site distinct from that of NGF and causes TrkA and ERK1/2 phosphorylation (Clewes et al., 2008). These distinct functions urge the necessity of getting more functional and structural information about proNGF and the isolated pro-domain.

The structure of mature NGF was established relatively early on (McDonald et al., 1991), showing an obligate parallel dimer, with each of the protomers forming a beta-sandwich. At contrast, little is known about the structure of proNGF. Preliminary far-UV circular dichroism (CD) data showed that the isolated pro-domain is monomeric and largely disordered with some evidence of secondary structure (Kliemannel et al., 2004). Attempts to solve the proNGF structure by NMR methods have so far failed due to the spectral complexity which not only reflects the size of the proNGF homodimer (50 kDa) but is also the probable consequence of a conformational equilibrium in the intermediate time-scale (Paoletti et al., 2011). A crystal structure of a proNGF complex with p75^NTR^, with a symmetric binding stoichiometry of 1:1, was determined at 3.75 Å resolution (Feng et al., 2010). However, the region of the electron density map corresponding to the pro-domain of proNGF could not be traced. Only sparse fragments of electron density were observed in the asymmetric unit that could not be easily connected. The absence of a defined trace of the proNGF pro-domain was thus interpreted as the consequence of flexibility, as also supported by our previous solution SAXS and NMR studies (Paoletti et al., 2009, 2011).

The presence of interactions between the pro-domain and NGF are supported by various functional and biochemical hints. The covalently attached pro-domain acts as an intramolecular chaperone since its presence significantly increases the yield and rate of *in vitro* refolding as compared to those of mature NGF (Rattenholl et al., 2001). In the crystal structure of the proNGF - p75^NTR^ complex (Feng et al., 2010), loops II of the mature NGF dimer have a conformation different from that observed for mature NGF in other complexes (He and Garcia, 2004; Wehrman et al., 2007) and in the unligated NGF (Holland et al., 1994; McDonald et al., 1991) suggesting possible interactions with the proNGF pro-domain. Chemical denaturation of proNGF yields two distinct transitions which likely correspond to disruption of the interacting surface between the pro-domain and mature NGF, resulting in the unfolding of the latter (Paoletti et al., 2011). Fluorescence and H/D exchange measurements indicated contacts between W142 of the NGF domain with residues W37-A57 of the proNGF pro-domain (Kliemannel et al., 2007). It was also recently reported that the pro-domain in proNGF induces a structural stabilization of loops I, II and IV in the NGF part, suggesting a direct interaction between residues R81-F89 of the proNGF pro-domain and loops I, II and IV in the NGF part (Trabjerg et al., 2017).

Here, we undertook an alternative strategy to prompt the functional role and acquire new structural information using a recombinant construct spanning the sequence of the mouse pro-domain (NGFpd), both in isolation and in the context of proNGF. Using a combination of advanced NMR experiments, we prove that the isolated NGFpd is able to interact, albeit weakly, with NGF. Distance restraints mapped by paramagnetic relaxation enhancement were then used in combination with fully atomistic Molecular Dynamics simulations to provide experimentally based structural models of proNGF which could be validated against previous SAXS data (Paoletti et al., 2011). We also demonstrate that NGFpd has *per se* the ability to induce growth cone collapse indicating a functional independence of this region and report direct evidence that the pro-domain is unfolded with well-defined tracts of local secondary structure. Our data thus represent an important step towards the structural and biochemical understanding of the properties of proNGF and the pro-domain.

## Materials and Methods

### Protein production

Expression and purification of NGFpd were carried out under native conditions. The expression plasmid for mouse NGFpd corresponding to residues 19-121 of mouse β-proNGF (UniProtKB P01139) (**Figure 1**) was prepared by truncation of pET11-proNGF expression plasmid used previously (Paoletti et al., 2011). The expressed protein contained an additional N-terminal methionine. Cloning and amino acid substitutions to introduce either E19 to Cys (NGFpdE19C), S24 to Cys (NGFpdS24C) or S90 to Cys (NGFpdS90C) mutations into NGFpd expression vector were made by inverse PCR and blunt end ligation. Expression of the unlabelled NGF and proNGF was carried out, as previously described (Paoletti et al., 2009; Rattenholl et al., 2001), in Luria Broth (LB) medium whereas M9 medium with the relevant isotopic enrichment was used for expressing isotopically ^15^N-labelled proteins for NMR. NGFpd mutants were expressed and purified as for the wild-type. ^15^N and ^2^H doubly labelled proNGF was expressed, refolded and purified according to a previous protocol (Paoletti et al., 2016), but growing the cells in perdeuterated water.

**Figure 1.**
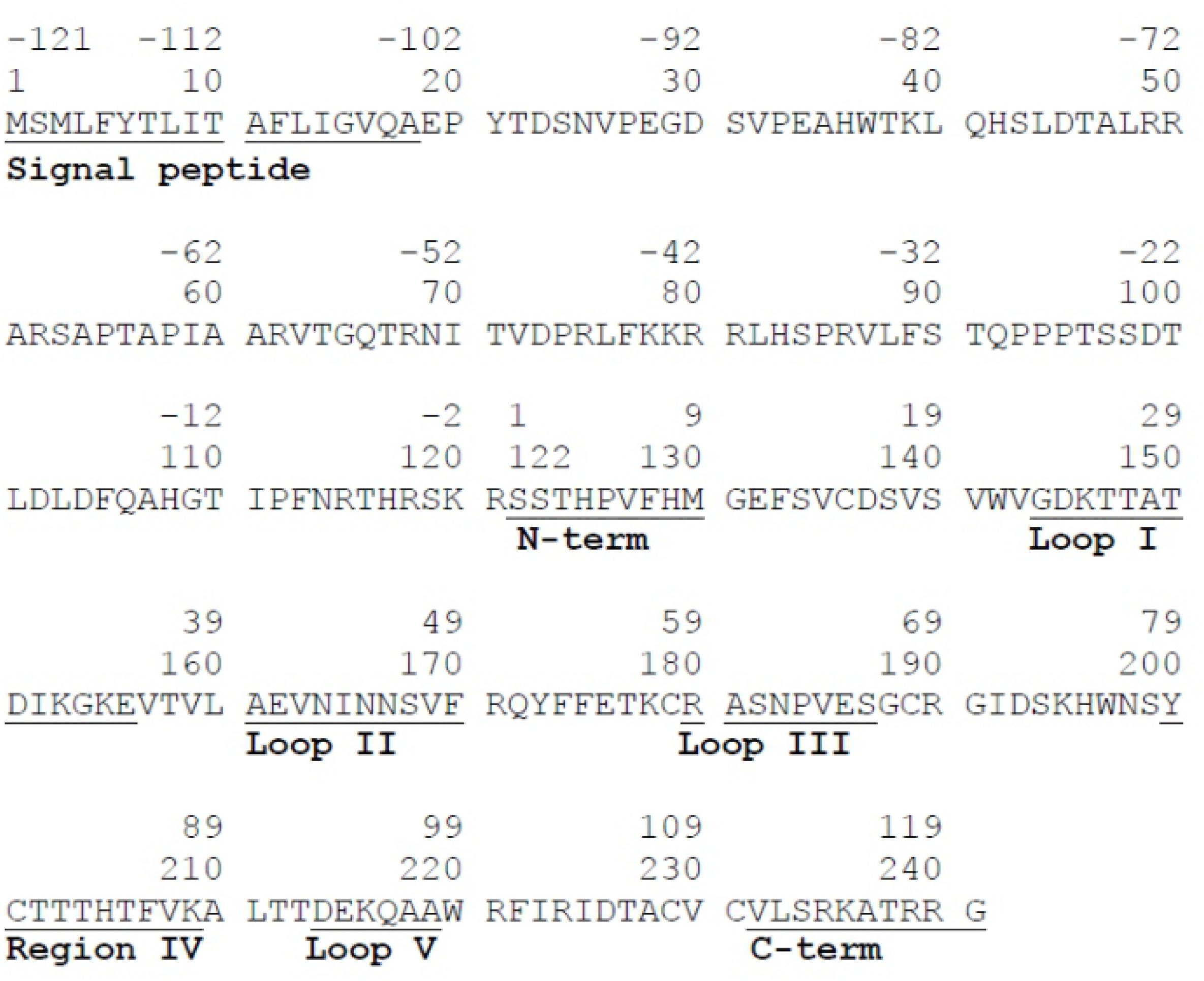
Primary structure of mouse proNGF. The amino acid numbering follows the convention in Rattenholl *et al*., 2001 (top line) and the UniProtKB accession number P01139 (bottom line). Underlined residue intervals are labeled according to Ibáñez (1995).

### Growth Cone Collapse Assays

Mouse hippocampal neuron cultures were prepared from P0 B6129 mice as described (Gobbo et al., 2017). At div 0 (day in vitro 0), 25-50,000 cells/cm^2^ were seeded on poly-D-lysine coated glass coverslips and grown at 37°C under 5% CO2 humidified atmosphere. On div 1 neurons were transfected with plasmids encoding P75^NTR^-GFP (Marchetti et al., 2014) or soluble GFP (pEGFP-N1, Clontech) along with pTagRFP-actin (Evrogen) with Lipofectamine2000 (Invitrogen). After 2 h, medium was changed to Neurobasal A (Gibco) supplemented with 2% B27 (Invitrogen), 2 mM glutamine (Invitrogen), 10μg/ml gentamicin (Invitrogen), 12.5 μM glutamate (Sigma-Aldrich) (culture medium). On div 3, neurons were incubated for 30 min with various neurotrophin forms or culture medium alone (untreated samples). Cultures were then fixed in 4% PFA 5% sucrose in PBS at room temperature for 15 min, washed twice in PBS, once in ddH2O, dried and mounted in Fluoroshield with DAPI (Sigma-Aldrich). The proNGF effect was tested at 10 ng/ml (0.4 nM), 100 ng/ml (4.0 nM) and 1000 ng/ml (40 nM). NGF and NGFpd were incubated at the same molar concentration (0.4, 4.0 and 40 nM) (molarity expressed as the concentration of the monomeric species). Cultures were imaged by confocal microscopy (Leica TCS SP5 on DM6000, equipped with MSD module) using an oil objective HCX PL APO CS 40.0X (NA=1.25). Sequential illumination with DPSS 561 and Ar 488 laser lines was used for TagRFP-actin and GFP imaging. Analysis was performed with ImageJ (available at NIH). Growth cones were identified by morphology and accumulation of fluorescent actin.

### CD Measurements of NGFpd

Far-UV CD measurements were carried out on NGFpd and its mutants at a concentration of 113 μg/ml in 50 mM sodium phosphate, 50 mM NaF pH 6.8 at 25°C using a 1 mm cuvette on a Jasco J-1100. The HT voltage was below 600 V over the entire scan range of 260-190 nm. Baseline corrected data were analysed using the CONTIN/LL method and reference set 4 on the Dichroweb server (Provencher and Glöckner, 1981; Sreerama and Woody, 2000; Whitmore and Wallace, 2008).

### NMR backbone assignment and secondary structure characterisation of NGFpd

^15^N-HSQC, HNCACB, CBCA(CO)NH, HN(CA)CO and HNCO experiments (Salzmann et al., 1999) were recorded on 800 μM [^15^N, ^13^C]-uniformly labelled NGFpd in 50 mM sodium phosphate at pH 6.8, 50 mM NaCl and 1 mM EDTA at 25°C on an AvanceIII Bruker 600 MHz spectrometer equipped with a TXI cryoprobe. Data were processed with nmrPipe (Delaglio et al., 1995) and analysed using CCPN Analysis (Vranken et al., 2005). Backbone resonances were assigned for 86 over 93 non-prolyl residues, equating to ∼92% of the assignable backbone resonances. The missing assignments of NGFpd include the first two N-terminal residues H42 and S43 and residues H117-S119 adjacent to the C-terminus. The C^α^, C^β^ and C’ shifts were used to calculate the chemical shift index (CSI) (Wishart and Sykes, 1994) using nmrView (Johnson and Blevins, 1994). The secondary structure propensity relative to the C^α^ and C^β^ shift were calculated using the SSP algorithm (Marsh et al., 2006).

### Paramagnetic Relaxation Enhancement Data Collection and Analysis

(1-Oxyl-2,2,5,5-tetramethylpyrroline-3-methyl)methanethiosulfonate (MTSL) was dissolved in DMSO. To generate MTSL labelled NGFpd cysteine mutants (NGFpdE19C-MTSL, NGFpdS24C-MTSL and NGFpdS90C-MTSL), NGFpdE19C, NGFpdS24C and NGFpdS90C were each incubated with a 1000 fold molar excess of DTT for 10 minutes. DTT was removed by passing each sample through a PD-10 column. The MTSL-DMSO solution was subsequently added to give a 30 fold molar excess of MTSL to protein, and a final DMSO concentration of 10% (v/v). Excess MTSL was removed after 2 hours from each sample by passing the sample through a PD-10 column. The samples were concentrated and dialysed into 50 mM sodium phosphate at pH 6.8, 50 mM NaCl and 1 mM EDTA.

^15^N HSQC spectra were recorded for ^15^N labelled NGFpdE19C-MTSL, NGFpdS24C-MTSL and NGFpdS90C-MTSL. Paramagnetic and diamagnetic spectra (*i.e.* in which the MTSL label was in the paramagnetic or diamagnetic states) were recorded in the absence and in the presence of L-ascorbic acid. Paramagnetic and diamagnetic ^15^N HSQC spectra were also recorded for both ^15^N labelled NGF and ^15^N labelled NGFpd (WT) in the presence of 0.5 molar equivalence of ^14^N labelled NGFpdE19C-MTSL, NGFpdS24C-MTSL or NGFpdS90C-MTSL. Data were treated as described (Battiste and Wagner, 2000) to calibrate distance restraints between the MTSL label and the backbone and sidechain NH groups from the paramagnetic to diamagnetic intensity ratios (I^para^/I^dia^).

### Structural modelling, simulations and refinement

A Flexible-meccano algorithm that efficiently generates ensembles of molecules, on the basis of amino acid-specific conformational potentials and volume exclusion and in particular amenable to intrinsically disordered proteins, was used to generate a structural ensemble of NGFpd (Ozenne et al., 2012). Additional structural propensities were included in the calculation based on the Secondary Structure Propensity (SSP) analysis (Marsh et al., 2006). These included helical propensities in the regions E34-R52, I59-R62, R75-K78, T96-T110 and N114-T116. In addition, two long-range contacts between two different regions of NGFpd were included as interpreted from intramolecular-PRE measurements: we imposed average distances of 16.5 Å between A35-W37 and F89-T91 and 14.5 Å between D23-N25 and T71-R75. A total of 10,000 conformers were calculated. Of these conformers, two representative structures significantly differing in helical content and radius of gyration were used as the starting point for fully solvated molecular dynamics simulations (MD). To reduce the computational cost and since NMR data indicated no significant interaction between individual pro-domains, only one pro-domain was attached to the mature NGF dimer (PDB ID: 1BET) using Modeller 9.12 (Fiser and Sali, 2003), leading to one proNGF chain with the second chain containing only mature NGF. Two unrestrained MD simulations were performed for 3 μs at 298 K in the NVT ensemble using the GROMACS software with the AMBER 99SB* ILDN protein force-field, and the dispersion-optimised TIP4PD water model. This force-field was recently shown to better capture the ensemble feature of disordered or partially disordered models (Piana et al., 2015). Na+ and Cl-ions (100 mM) were added to the solution to mimic the experimental conditions. From the full trajectory, 30,000 structures were extracted (every 100 ps) and analysed. The complete proNGF dimer was generated by combining 57 representative structures of the most populated conformers of NGFpd into 1,596 different dimer structures. Structures with steric clashes among the NGFpd parts were removed from the ensemble leaving 1,486 structures. The relative population of the unique dimeric structures was obtained by combining the relative population of each monomeric conformer. The subsequent analyses and comparison to experiments were carried out on these structures. The radius of gyrations was calculated using CRYSOL. The distances corresponding to the NMR derived contacts were computed with the Python MDTraj library.

### SAXS Validation

Analysis of the overall parameters was carried out by the PRIMUS software suite (Konarev et al., 2003) from ATSAS package (Franke et al., 2017). Inter-domain flexibility and size distribution of possible conformers of proNGF in buffer were quantitatively assessed by the ensemble optimization method (EOM) (Bernado et.al., 2007). In EOM, the pool of about 2000 conformers derived from MD simulations was analysed. The theoretical scattering pattern was calculated for each generated model by CRYSOL (Svergun et al., 1995). A genetic algorithm (GAJOE) was used to select an ensemble of conformations whose mixture best fitted the experimental data. Multiple runs of EOM were performed and the obtained subsets analyzed to yield the distribution of the radius of gyration (Rg) in the selected ensembles. Once each ensemble was determined, the corresponding Shannon Entropy, reported as Rflex, provided a quantitative measure of flexibility (Tria et.al. 2015).

## Results

### NGFpd is an intrinsically disordered protein with helical secondary structure propensity

The CD spectrum of NGFpd showed a minimum at 201 nm consistent with a largely disordered protein with some elements of secondary structure (**Figure S1**) in agreement with previous data (Kliemannel et al., 2004). Deconvolution of the spectrum suggested a secondary structure content of 20% α-helix and 20% β-sheet.

The ^15^N-HSQC of NGFpd had limited dispersion in the proton frequency of the backbone amide resonances (7.8-8.5 ppm) (**Figure S2**). Despite this, the resonances were fairly well resolved which greatly facilitated backbone resonance assignments. The single tryptophan indole NH^ε^ of W37 was clearly visible at 10.1 ppm. The ^15^N NOESY-HSQC of NGFpd contained limited NOEs, with no high-field shifted signals for aliphatic residues below 0.7 ppm. The CSI of NGFpd indicated secondary structure propensity (**Figure 2A**) with several shifts towards a helical conformation between P33 and R49 and β-sheet conformation between T67 and D73. The SSP plot showed a significant stretch, spanning residues E34 to R49, with a 26-45% α-helical propensity (**Figure 2B**). A shorter stretch, containing residues R75 to K78, indicated a more limited α-helical propensity.

**Figure 2.**
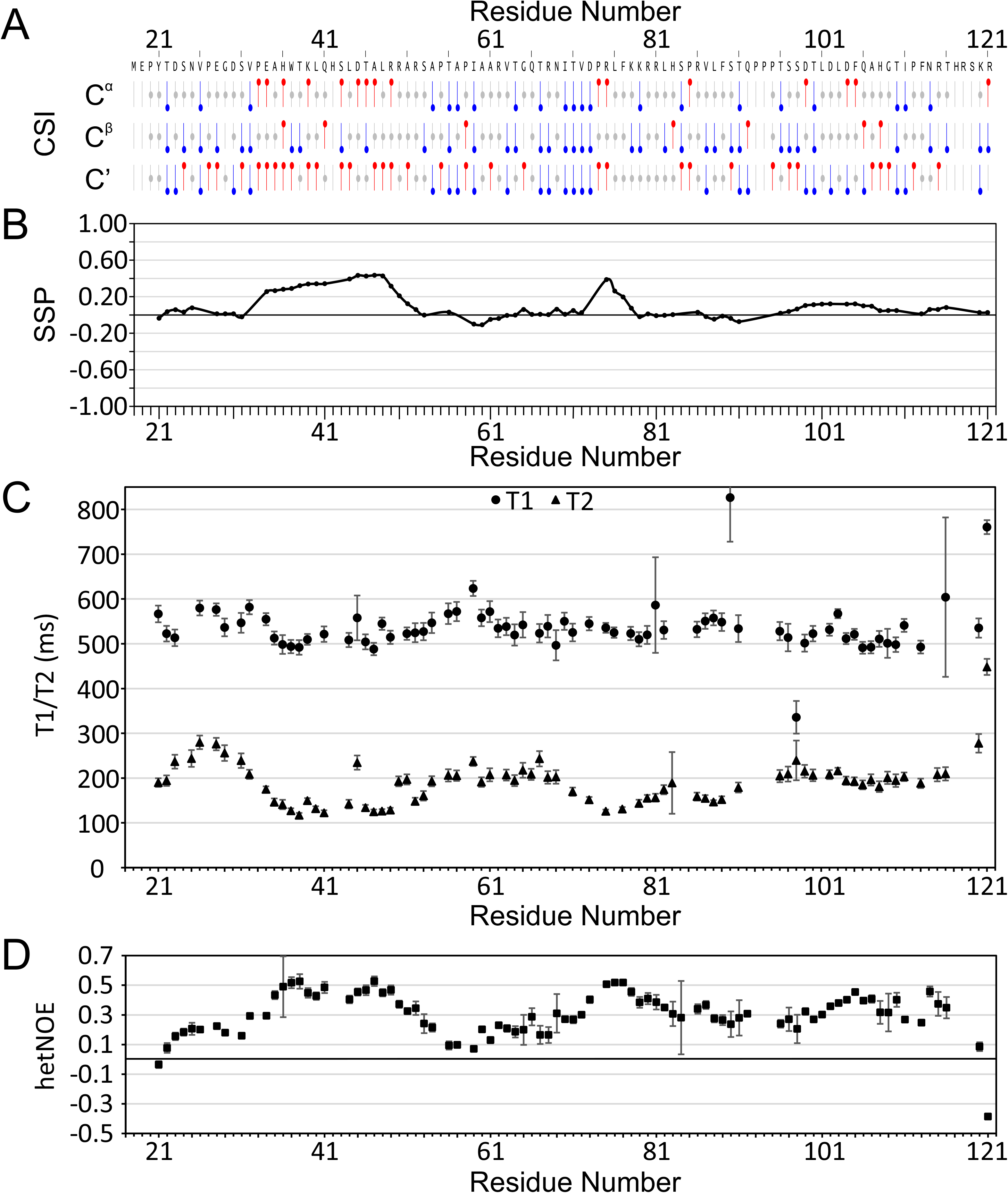
Secondary Structure Propensity of NGFpd. **A) C^α^, C^β^ and C’ secondary shifts**, as visualised using NMRViewJ. Red “lollipops” indicate α-helical shifts, blue “lollipops” indicating β-sheet shifts. **B**) Secondary Structure Propensity (SSP) analysis of NGFpd, where a value of +1 indicates 100% α-helical and −1 indicates 100% β-sheet. **C**) T1 and T2 values for NGFpd at 800 MHz. **D**) Heteronuclear-NOE values for NGFpd.

To assess the dynamics of NGFpd, we recorded T1, T2 and ^1^H-^15^N heteronuclear NOE measurements (**Figure 2C,D**). The average T1 and T2 values are 530 ms and 190 ms respectively. Deviations from the average, indicating reduced flexibility, were observed for residues 35-49, whose average T1 and T2 values are 510 ms and 140 ms respectively, and for residues 75-78, whose T1 and T2 values were 525 and 135 ms respectively. The average ^1^H- ^15^N heteronuclear NOE value is 0.31, with deviations towards less dynamic conformations for residues 35-49 and 75-78, whose average values are 0.47 and 0.5 respectively.These regions correlate with the residues predicted to have secondary structure propensity.

These results confirmed previous studies (Kliemannel et al., 2004; Paoletti et al., 2011) and mapped the regions of NGFpd which retain local secondary structure.

### The NGFpd has *per se* the ability to induce growth cone collapse

We then assessed whether NGFpd could *per se* retain functions observed for uncleaved proNGF and tested this hypothesis by carrying out a growth cone collapse assay, the best validated assay for proNGF function (Deinhardt et al., 2011). We observed that proNGF caused growth cones collapse in p75^NTR^-overexpressing mouse hippocampal neurons (**Figure 3A,B**) as previously reported (Deinhardt et al., 2011). Surprisingly though, we found that also the NGFpd alone was capable of collapsing growth cones, whereas NGF was not (**Figure 3A**). The mechanism of NGFpd action is likely depending on its binding to p75^NTR^, in analogy to proNGF. The effect of NGFpd (as well as of proNGF) was smaller in neurons transfected with GFP alone (**Figure 3B,C**). This is consistent with the observation that a subpopulation of hippocampal neurons expresses p75^NTR^ and is responsive to proNGF (Deinhardt et al., 2011). Accordingly, neurons transfected with p75^NTR^ were responsive to proNGF as well as to NGFpd treatment in a dose-dependent manner (**Figure 3C**). The effect is not due to p75^NTR^ expression on its own, since untreated neurons display a comparable number of intact growth cones to control neurons (**Figure 3C,D**). We therefore concluded that NGFpd is as effective as proNGF in causing growth cones to collapse (**Figure 3C,E**) and is thus able to carry this function independently.

**Figure 3.**
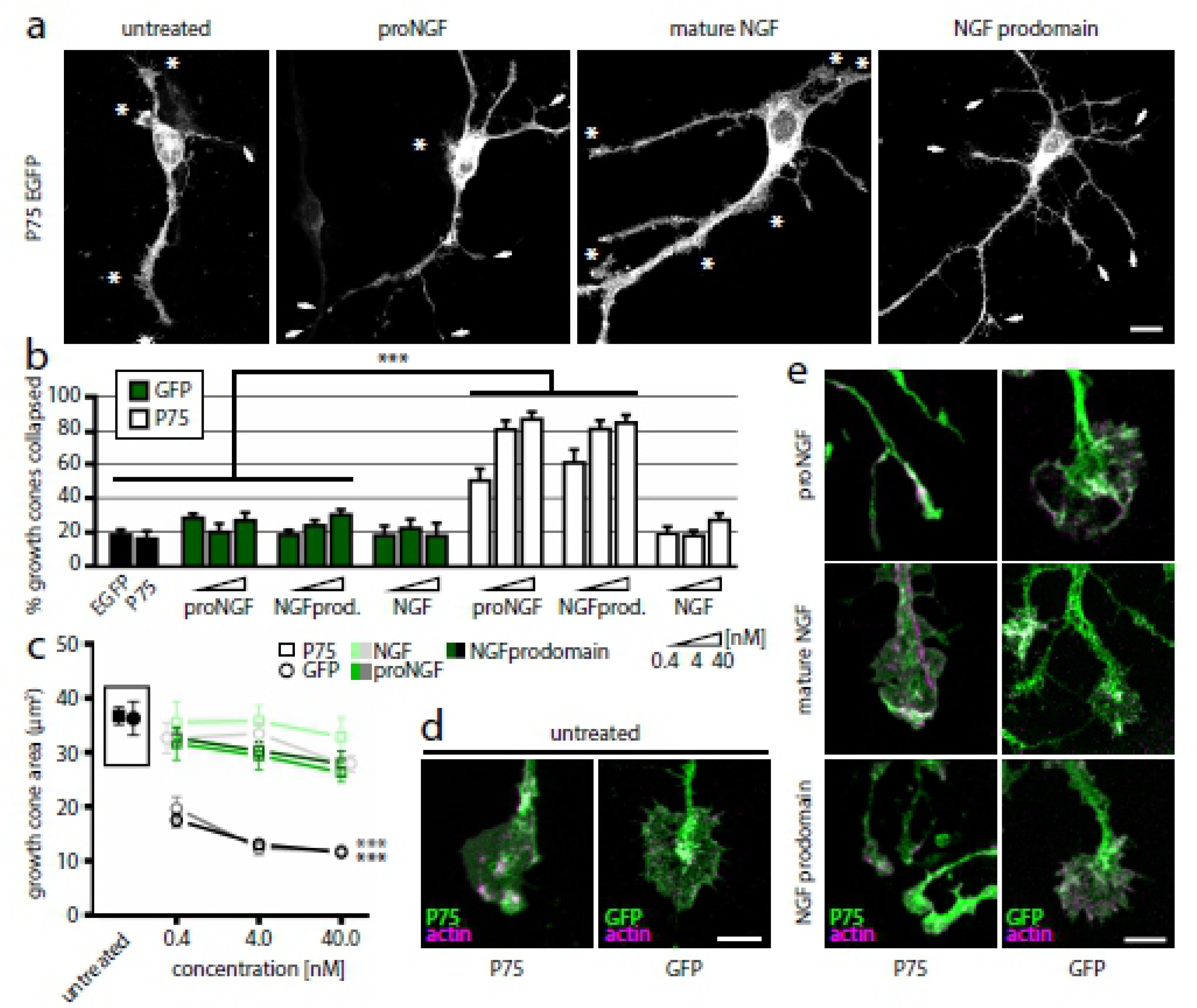
NGFpd is capable of inducing growth cone collapse. **A**) Treatment of div3 neurons expressing P75^NTR^ with 4 nM proNGF (which corresponds to 100 ng/ml proNGF) for 30’ induces growth cones collapse. In untreated neurons, the majority of growth cones remain intact (asterisks); collapsed cones are indicated with arrowheads. 4 nM NGFpd, but not 4 nM mature NGF, induces growth cone collapse. **B**) Fraction of collapsed growth cones in neurons transfected with p75^NTR^-EGFP (white bars) or EGFP (green bars), and treated with 0.4, 4 or 40 nM of each neurotrophin, or untreated (black bars). Each set of three represents, left to right: 0.4 nM, 4 nM and 40 nM. p75^NTR^-GFP samples display a dose-dependent increase in the fraction of collapsed growth cones when treated with proNGF or NGFpd, but not with NGF. The effect was minor, but still observed, in neurons transfected with GFP only. *** P<0.001, pairwise multiple comparison, Kruskal-Wallis test. n=8-56 from 2-4 independent samples for each sample. **C**) Growth cone area after 30’ treatment. Circles, neurons expressing p75^NTR^-GFP; squares, neurons expressing GFP; filled symbols are from untreated cultures. p75^NTR^-GFP, NGF treated symbols (grey circles) at 0.4 nM and 40 nM are moved slightly aside for clarity of presentation. *** P<0.001 compared to NGF, NGFpd vs. proNGF P=0.89; two-way ANOVA, Tukey test (factor A treatment df = 2, F=141.6; factor B concentration df=2, F=12.68). n=51-267 from 2-4 independent samples for each point. **D**) Representative growth cones of p75^NTR^-GFP and GFP neurons. p75^NTR^ does not determine growth cone collapse in untreated neurons. (**E**) Growth cones from p75^NTR^-GFP (left) and GFP (right) neurons after 30’ treatment with 0.4 nM proNGF, mature NGF or NGFpd. RFP-actin is in magenta, GFP in green. Data are means±SEM. Scale bar (a) 10 μm (d,e) 5 μm.

### NGFpd interacts in trans with mature NGF

We next tested whether isolated NGFpd could interact with NGF. We recorded a ^15^N TROSY HSQC spectrum of deuterated ^15^N NGFpd titrated with up to 5 molecular equivalents of unlabelled NGF monomer (i.e, 2.5 molecular equivalents of NGF dimer) (**Figure S3A**). We observed differential peak broadening and some fast exchange resonance shifts, indicating an interaction. Since line broadening is quite severe above 1.5 molecular monomer equivalents (0.75 NGF dimer equivalents), we analysed the chemical shift perturbations (CSP) at 1.5 molecular equivalents (**Figure 4A**). Shifted resonances were observed also at 0.25 molecular equivalents (0.125 dimer equivalents) suggesting that the affinity is in the μM-mM range. The largest CSPs (greater than 0.02 ppm) were observed for Y21, V26, R49, F77 and T116. CSPs greater than or equal to 0.01 ppm were observed for several other resonances. In addition, the resonances of V87, L88, F89, S90 and T91 showed significant line broadening that prevented observation of their CSPs already at 1.5 molecular equivalents (0.75 dimer equivalents). Severe line broadening at 0.2 molecular equivalents was observed for T91, whereas V87 and L88 broadened only at 0.8 molecular equivalents. From this CSP analysis, we could thus identify two contiguous regions of NGFpd that exhibited the largest CSPs/line broadening around Y21-E28 and V87-T91.

**Figure 4.**
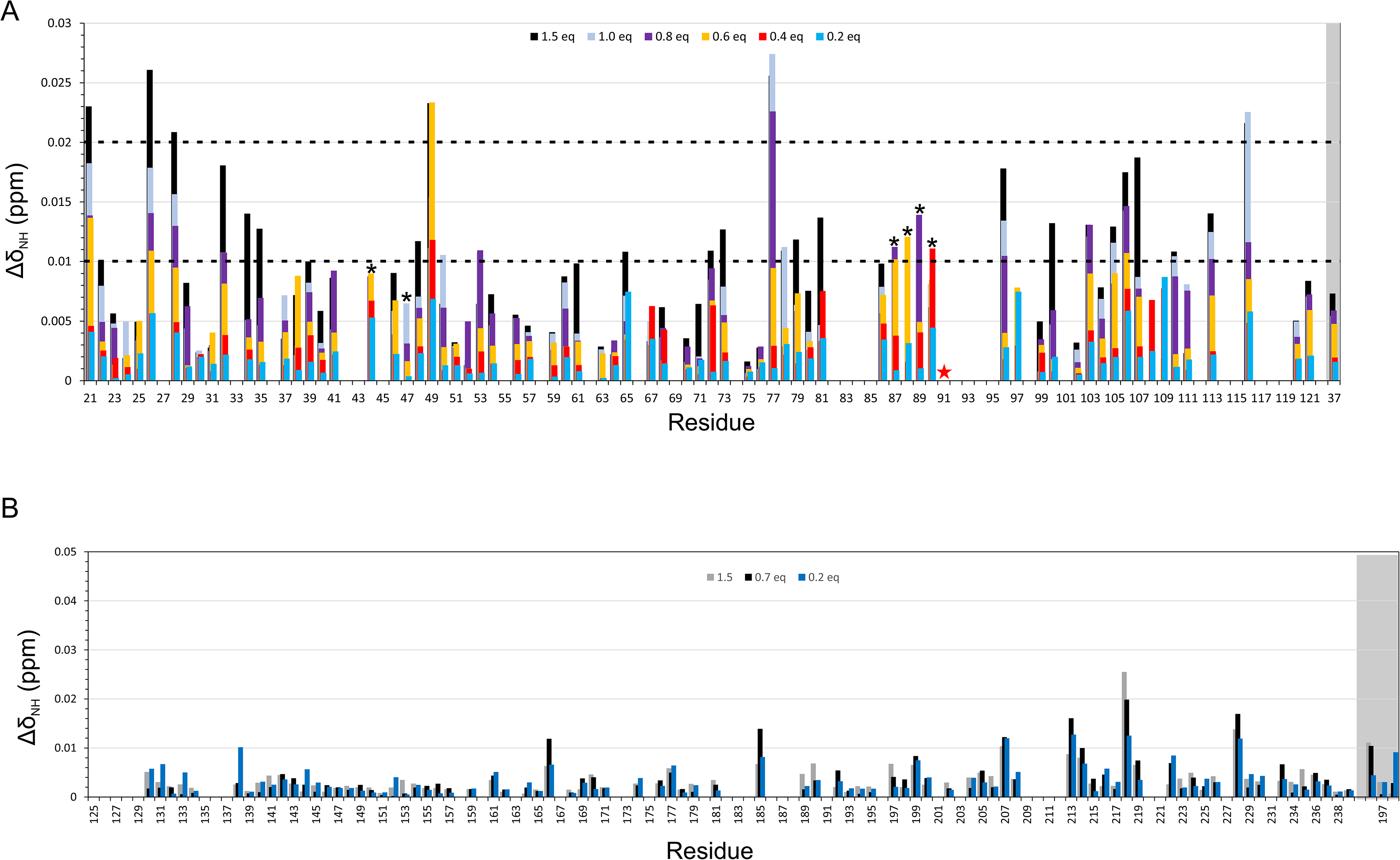
Effects of titrations of NGFpd with NGF and viceversa. (**A**) CSP per residue for the titration of ^15^N NGFpd with unlabelled NGF. The red star above residue 91 indicates severe line broadening at 0.2 equivalents and asterisks indicate the severe line broadening occurred between 0.8 and 1.5 equivalents (monomer:monomer ratio). The grey shaded box indicates the W73 indole NH. (**B**) CSP per residue for the titration of ^15^N NGF with unlabelled NGFpd. The grey shaded box indicates the tryptophan indole NH groups.

Vice versa, we titrated deuterated ^15^N NGF with up to 5 equivalents of unlabelled NGFpd (2.5 NGFpd monomer to NGF dimer) and recorded ^15^N TROSY HSQC spectra for each ratio (**Figure S3B**). We observed severe differential line broadening effects, confirming an interaction between the two. The magnitudes of the CSPs are very small (**Figure 4B**). The largest CSPs were observed for residues N166 (Loop II), V185 (Loop III), T213, A218 (Loop V) and A228 and the indole of W142. The greatest increases in linewidth were observed for residues T177, C179, A219 (Loop V), W220 and R221, whereas the greatest reductions in intensity were for residues A161 (Loop II), E162 (Loop II), F170 (Loop II), T177, D214 (Loop V) and the indoles of W142 and W220.

### NGFpd forms intramolecular tertiary contacts

To obtain indications on the conformational tendencies of the NGFpd random coil we used Paramagnetic Relaxation Enhancement (PRE), since this technique has proven to be effective in measuring transient long distance contacts that occur in intrinsically unfolded proteins (Clore et., 2007). In this experiment, a paramagnetic probe is attached to cysteines. Residues in spatial proximity of the label will broaden in the spectrum. We used the nitroxide spin label MTSL, individually attached covalently *via* a thioester bond to residues of NGFpd (E19C, S24C or S90C), that were mutated to a cysteine (wild-type NGFpd has no cysteines). These positions were chosen because they are adjacent to the regions of NGFpd where the most significant changes were observed in the titrations of NGFpd with NGF: we were of course wary of introducing mutations that might prevent interaction, which is why we chose to mutate residues that were adjacent to the greatest CSPs rather than mutate the residues exhibiting the greatest CSPs. The HSQC spectra of the mutants confirmed that the mutants remain unstructured and monodisperse (**Figure S4**). The information was then detected by measuring the paramagnetic/diamagnetic ratios of the ^15^N HSQC spectra (**Figure 5 and S4**). Extreme peak broadening (I^para^/I^dia^ < 0.1) was observed up to 5 residues away from the MTSL label for NGFpdS24C-MTSL (**Figure 5B**), followed by a gradual diminishing PRE effect for residues more sequentially distant from the probe. Significant broadening was also observed for D73 (I^para^/I^dia^ 0.25) and S84 (I^para^/I^dia^ 0.15), which are not sequentially close to the probe, indicating that the MTSL probe must be close in space to these residues. Similarly, extreme PRE effects were observed for residues of the NGFpdS90C-MTSL mutant (**Figure 5C**) which are sequentially close to the probe, but also for residues 32-44 and 69-73, again indicating that the probe is close in space to these residues.

**Figure 5.**
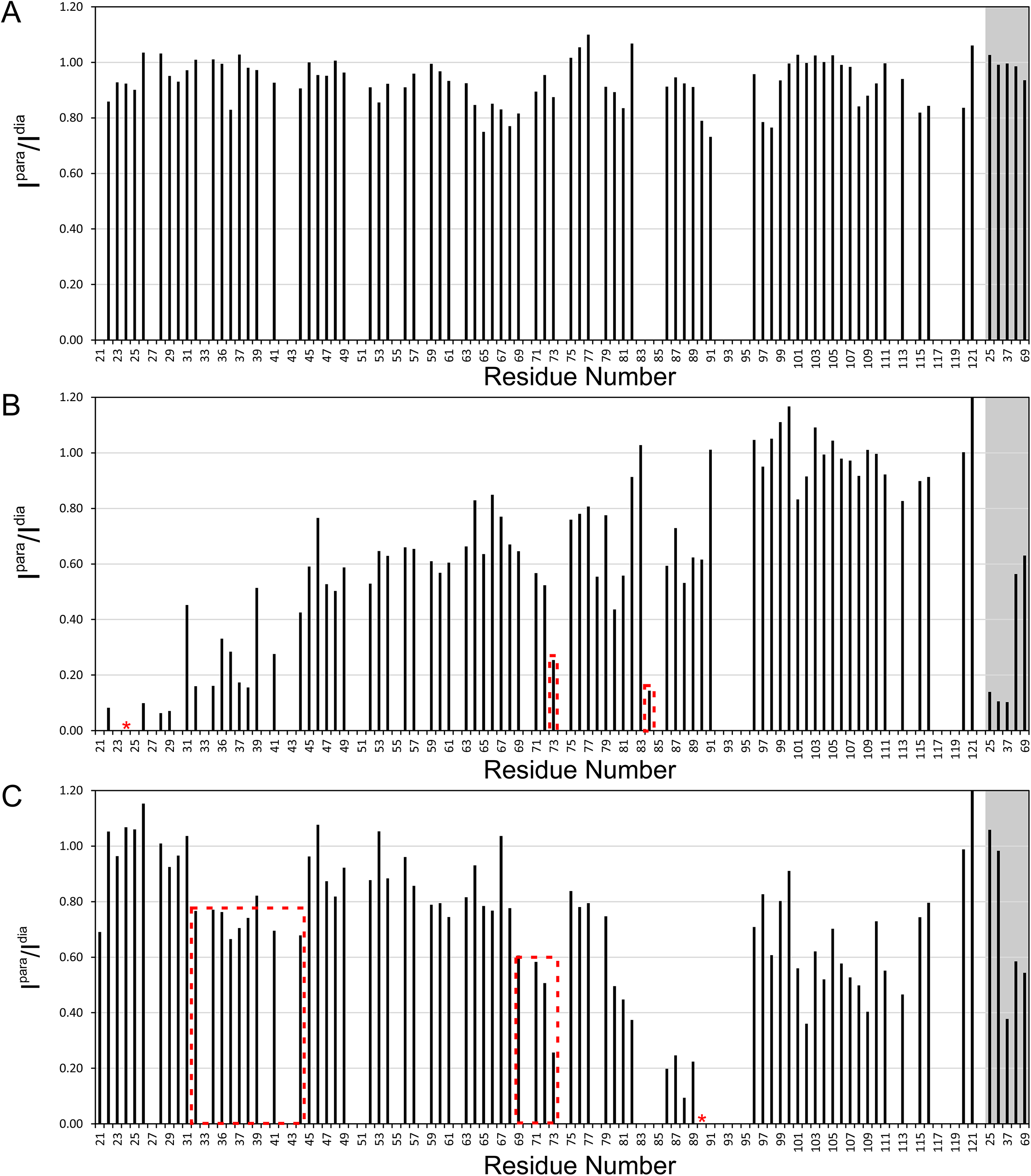
Normalised paramagnetic/diamagnetic intensity ratios (I^para^/I^dia^) per residue for MTSL labelled NGF pro-domain mutants. **A**) ^15^N labelled NGFpdE19C-MTSL, **B**) ^15^N labelled NGFpdS24C-MTSL, **C**) ^15^N labelled NGFpdS90C-MTSL. The red asterisks indicate the positions of the MTSL label and dotted red lines indicate regions of significant PRE. Sidechain NH groups are indicated by the grey box.

To ascertain whether the distant PRE effects were solely due to the formation of intramolecular tertiary contacts or caused by intermolecular contacts between two NGFpd molecules we recorded paramagnetic and diamagnetic HSQC spectra of ^15^N labelled NGFpd in the presence of ^14^N labelled NGFpdE19C-MTSL, NGFpdS24C-MTSL or NGFpdS90C-MTSL (**Figure 5D,E,F**). If the effects were intermolecular we expected to see broadening of the ^15^N NMR signals of the ^15^N labelled NGFpd. No significant PRE effects were observed.

Taken together, these results indicate that NGFpd forms transient intramolecular tertiary contacts and has a clear three-dimensional propensity.

### PRE effects between NGFpd and NGF

To ascertain whether the shifts observed in the titration of labelled NGFpd with unlabelled NGF were caused by contacts with NGF or by conformational changes of NGFpd upon binding to NGF, we measured intermolecular PREs between the ^15^N labelled NGF and ^14^N labelled NGFpdE19C-MTSL, NGFpdS24C-MTSL or NGFpdS90C-MTSL. NGFpdE19C-MTSL and NGFpdS24C-MTSL would thus report on the Y21-E28 region of NGFpd, whereas NGFpdS90C-MTSL would report on V87-T91 region of NGFpd. In all three cases, intermolecular PREs were observed, suggesting that both the regions Y21-E28 and V87-T91 of NGFpd are located proximal to the NGF surface in the NGF-NGFpd interacting adduct. The PRE effects observed on NGF backbone amides by titration with NGFpdE19C-MTSL (**Figure 6A**) were moderate with the greatest effects observed for residues D23, S43, S97, R115 and H117 (I^para^/I^dia^ ratios <0.7-0.8), whereas significantly larger effects were observed for NGFpdS24C-MTSL (**Figure 6B**).

**Figure 6.**
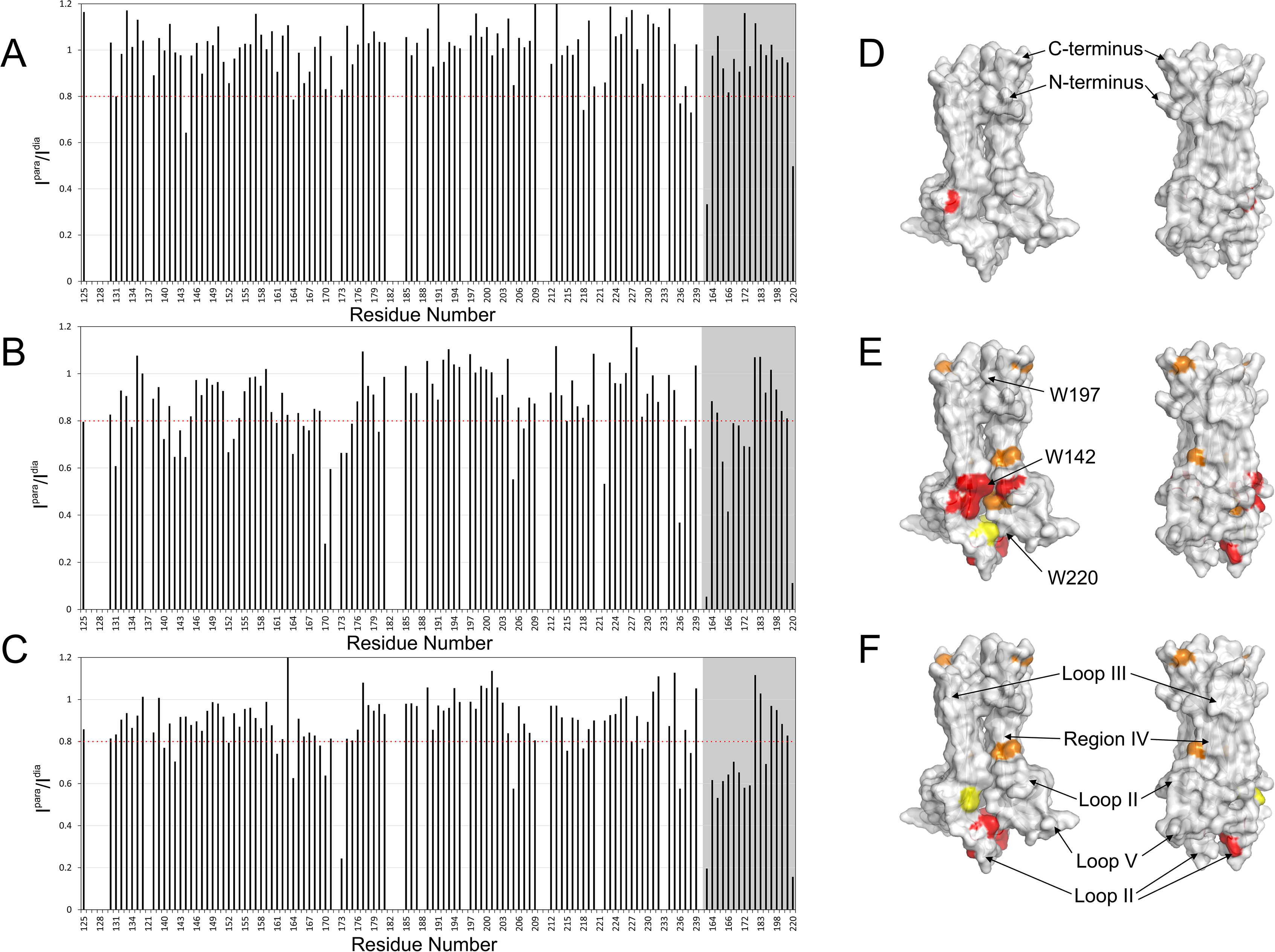
Normalised paramagnetic/diamagnetic intensity ratios (I^para^/I^dia^) per residue of mature ^15^N NGF in the presence of MTSL labelled ^14^N NGF pro-domain mutants. **A**) ^15^N labelled NGF with ^14^N labelled NGFpdE19C-MTSL (1:0.5 mix, monomer:monomer). **B**) ^15^N labelled NGF with ^14^N labelled NGFpdS24C-MTSL (1:0.5 mix, monomer:monomer). **C**) ^15^N labelled NGF with ^14^N labelled NGFpdS90C-MTSL (1:0.5 mix, monomer:monomer). Significant PREs are indicated below the dotted red lines. Sidechain NH groups are indicated by the grey box. I^para^/I^dia^ ratios of less than 0.3, 0.6 and 0.7 are mapped onto the surface of NGF (PDB ID:1BET) in yellow, orange and red respectively for **D**) ^15^N labelled NGF with ^14^N labelled NGFpdE19C-MTSL (1:0.5 mix), **E**) ^15^N labelled NGF with ^14^N labelled NGFpdS24C-MTSL (1:0.5 mix) and **F**) ^15^N labelled NGF with ^14^N labelled NGFpdS90C-MTSL (1:0.5 mix).

Mapping these PRE effects onto the structure of NGF localised these regions adjacent to the dimer interface, approximately between W142 and W220 (**Figure 6D, E, F**). The effect was stronger with NGFpdS24C-MTSL, suggesting that residues Y21-E28 of NGFpd are involved in binding to this region of NGF. The effects for NGFpdS90C-MTSL (**Figure 6C**) were milder but still broadly localised in the same area, suggesting that residues V87-T91 are also close to this binding surface but not as close as Y21-E28. Since long range intramolecular PREs were observed for NGFpdS90C-MTSL at residues V32-E34 of NGFpd, which are sequentially adjacent to Y21-E28, we considered conceivable that the tracts Y21-E28 and V87-T91 of NGFpd could be both located near the same binding surface of NGF. Also, residues Y21-E28 should be located at the NGF binding site identified by the PRE data, whereas residues V87-T91 should be close to this binding site by association with Y21-E28 given that the intermolecular PREs were stronger with NGFpdS24C-MTSL. The CSPs and the severe NMR signal line broadening observed for residues V87-T91 may therefore be the result of a conformational change in the NGFpd upon binding to NGF.

### Structural Modelling of NGFpd and proNGF

The combination of the information obtained from PRE and CSP resulted in 26 non-ambiguous inter-domain distance restraints with a mean and standard deviation of 16.9 Å and 1.6 Å, respectively. To reconstruct a model of proNGF, we first calculated a bundle of initial structures of NGFpd using the Flexible-meccano software (Ozenne et al., 2012). This process resulted in 10,000 structures, which showed a great variability in helical content and radius of gyration of NGFpd (0.99–3.07 nm, mean 1.87 and standard deviation 0.35 nm). In addition to the NMR restraints, we imposed helical restraints in the E34-R49 region. However, albeit the CSI data indicate helical conformations in ca. 40% conformers in the region E34-R49, the fractional helicity in the Flexible-meccano output ranged from 0 to 0.88% (mean 0.15%, standard deviation 0.31%).

From this bundle, two structures of NGFpd differing by radius of gyration (1.81 and 2.14 nm) and helical content (0.88 and 0%) were arbitrarily selected to build the starting structures of further refinement (**Figure 7A)**. Two partial NGFpd-NGF adducts were built using the different pro-domain models and used to run 3 μs fully solvated MD simulations (see Methods). After this time, the full dimer was reconstructed by combining 57 representative structures from the most populated clusters of NGFpd in the NGFpd-NGF adduct simulations. On average, trajectory B tended to satisfy all the experimental distances from S24, while trajectory A tended to satisfy the distances from S90 (**Figure S5A,B**), the satisfaction of distances from S24 and S90 being roughly mutually exclusive. Only a few short lived conformers satisfied NMR-derived distances from both sets but never more than 50% of each one of the sets. The Free Energy Surfaces (FES) from the two trajectories tended to visit different conformations (**Figure S5C,D).** Three regions were visited by both trajectories, labelled 0, 9 and 10 in trajectory A, and 0, 5 and 6 in trajectory B. The relative populations of the various conformers were estimated from the relative populations of the conformations visited by the two independent NGFpd-NGF adducts models (**Figure 7B**). The average helicity of the E34-R49 region was ∼24% with a standard deviation of 24%, improving the agreement with the experiments as compared to the Flexible-meccano ensemble. Of the two trajectories, trajectory A visits more helical conformers (average helicity 31% with a standard deviation of 28%). Interestingly, in both trajectories we frequently observe close contacts between residues R81-F89 of the proNGF pro-domain and loops I, II and IV in the NGF part (**Figure S5F**), in agreement with recent H/D exchange experiments (Trabjerg et al., 2017). A full dimer structure extracted from the simulations in which one pro domain satisfies >80% of the NGFpdS24C-MTSL contacts and the other >80% of the NGFpdS90C-MTSL contacts is shown in **Figure 7C**, while the E34-R49 region is in a helical conformation (its position is marked by a red cross in **Figure 7B**). The structure represents an energetically accessible, albeit scarcely populated, conformer.

**Figure 7.**
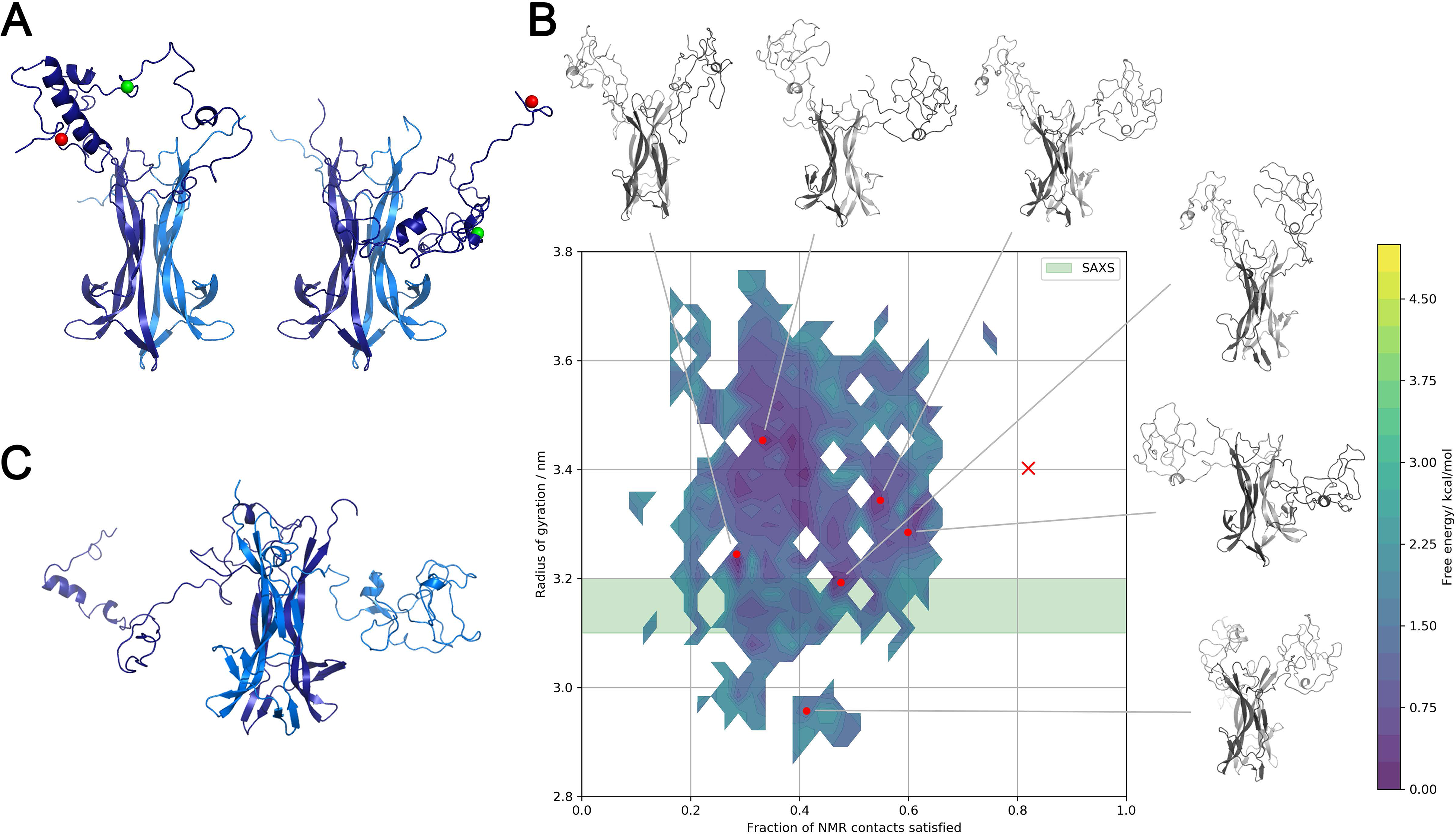
Molecular dynamics calculations of the proNGF dimer. **A**) Cartoon representation of the MD starting structures of the NGFpd-NGF dimer. The two conformations of the pro domain have been selected from the Flexible meccano structural ensemble. **B**) Approximated conformational free energy landscape of the NGFpd-NGFpd dimer as a function of the fraction of NMR contacts satisfied and the radius of gyration. The experimental radius of gyration is indicated by a green band. The fraction of satisfied NMR contacts was calculated using 26 unambiguous inter-domain distances, 16 from NGFpdS24C-MTSL and 10 from NGFpdS90C-MTSL. No ambiguous distances were included. Representative structures for the most populated clusters and for cluster satisfying most contacts are shown. **C**) Structure of the proNGF dimer that satisfies most NMR contacts and has a helical E34-R49 region (marked by a red x in panel B).

### Experimental validation by SAXS and NMR

We used previously recorded SAXS data on proNGF in 50 mM phosphate buffer (pH 7.0) (Paoletti et al., 2009) to validate the proNGF complex (**Figure 8A**, curve 1). Under these conditions, the estimated apparent molecular mass (MM_exp_ 45±5 kDa) and the hydrated particle volume (*Vp=*(*72*± 6)*10^3^ Å^3^) of proNGF agree with the presence of a proNGF dimer. The overall parameters (Rg = 31.5±0.5 Å, Dmax = 110±10 Å) point to the presence of more extended structures. EOM analysis of proNGF in buffer yielded high quality fit with χ^2^ value of 1.48 (**Figure 8A, curve 2**). Minor deviations at higher angles (*s* > 0.20 Å^−1^) can be explained by the presence of different conformations of NGFpd (Paoletti et al., 2016). The preponderant fraction of models in the optimized ensemble have Rg between 2.8-3.2 nm **(Figure 8A, insert).** A smaller fraction (with Rg between 3.3 and 3.4 nm) accounts for models with a more extended proNGF pro-domain supporting the hypothesis of considerable inter-domain flexibility. Quantification of the flexibility (ensemble Rflex = 60.6% versus pool Rflex = 83.4%) confirms numerically the flexibility of the pro-domain.

**Figure 8.**
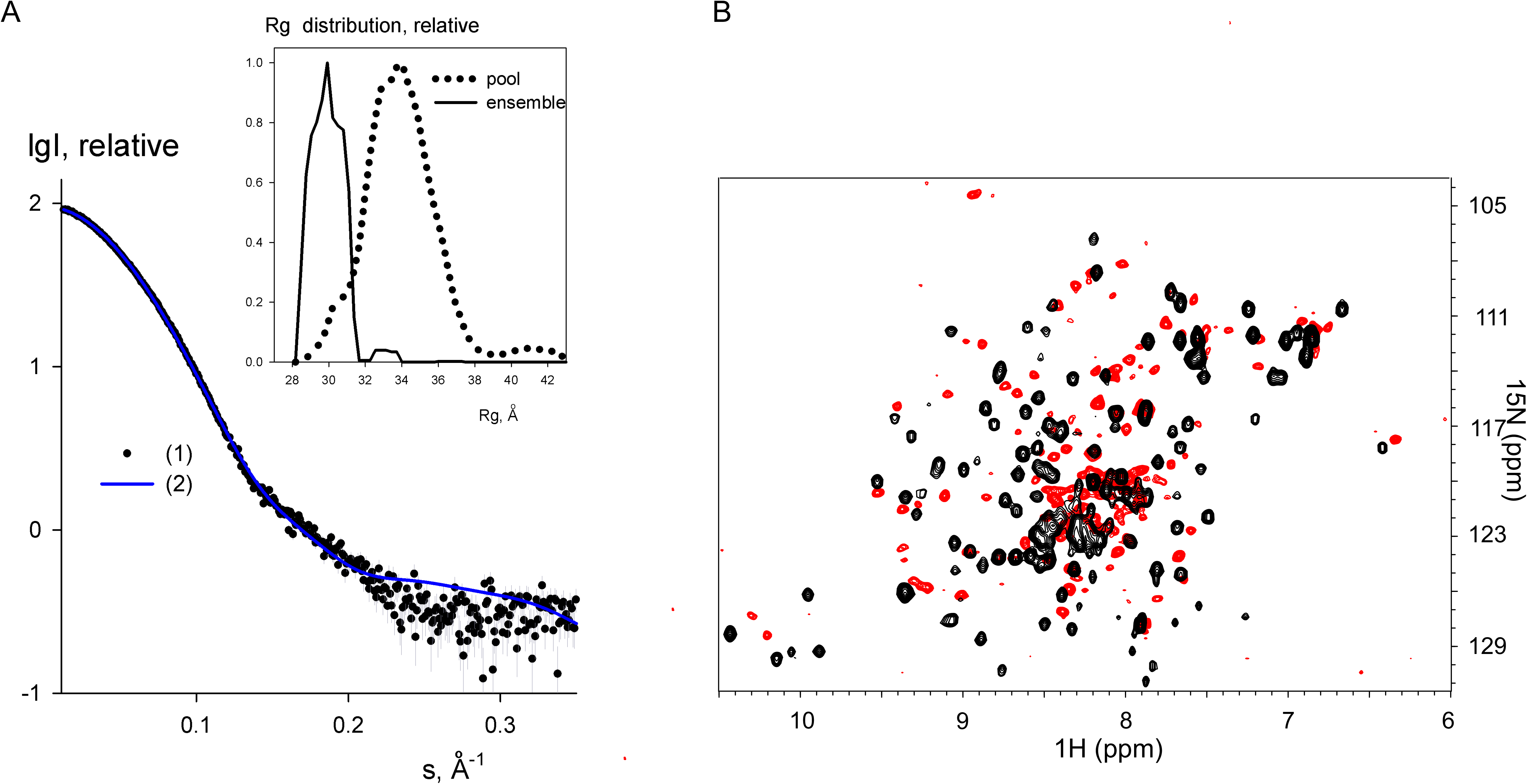
Validation of the model ensemble. **A)** Experimental SAXS data of proNGF in 50 mM Na-phosphate buffer (pH 7.0) are displayed as dots with error bars, the fit from the program EOM is indicated as blue solid line. R_g_ distributions of the EOM models for proNGF are displayed in the insert. The distribution of the initial pool of models derived from MD simulations are shown by dotted line, solid line corresponds to the selected ensembles. **B)** Comparison of the TROSY spectrum of proNGF and the HSQC of NGF recorded under similar conditions.

Independently, we also compared a high resolution ^15^N TROSY spectrum of proNGF with the HSQC of NGF to find support to our model. The overall features are very different (**Figure 8B**). Most notable are the positions of the tryptophan indoles which are shifted from those of NGF and the simplicity of the spectrum which would support a symmetric environment of the dimer. We also could identify a set of NMR resonances of NGF which are significantly shifted in the proNGF spectrum as compared to NGF. Their assignment was supported by a TROSY-modified NOESY on deuterated proNGF. The resonances correspond to residues (S122–M130) and (S234-T238), respectively at the N-and C-termini of NGF. Their behaviour is expected, being these stretches sequentially or spatially contiguous to the pro-domain in proNGF. A subset of resonances attributed to residues of mature NGF in or close to loops I (G144, T150, I152), II (F170) and V (T213, E215, A218, W220) had reduced intensities or were shifted. The resonances of residues belonging to loop III (A181–V189) were instead negligibly affected but had different NOE contacts. These findings confirmed the plasticity of the NGF loops (Paoletti et al., 2016) also in the context of proNGF. Finally, we noticed the absence in the proNGF spectrum of a set of resonances belonging to the central stem of the NGF domain (F174, F175, T177, V208, K209 and A210). This suggests a substantial change of chemical environment of the NGF domain within proNGF induced by the pro-domain, with residues R81-F89 causing a structural stabilization of loops I, II and V of NGF in agreement with recent HDX-MS findings (Trabjerg et al., 2017). Altogether the data are fully consistent with the experimentally based models.

## Discussion

Neurotrophins are key regulators of neuron survival and neurite extension and maintaining during development and adult life. Mature neurotrophins signal neurite extension predominantly *via* tyrosine-kinase receptors (Chao et al., 2006), while the unprocessed proneurotrophins, like proNGF and proBDNF, have the opposite function via a conserved mechanism that involves the neurotrophin receptor p75^NTR^ (Anastasia et al., 2015) and downstream Rac inactivation (Deinhardt et al., 2011; Sun et al., 2012). Treatment of central or peripheral neurons with sub-nanomolar concentrations of proNGF or proBDNF induces the rapid collapse of growth cones *via* actin depolymerization (Deinhardt et al., 2011). Increasing evidence also shows that the pro-domain of BDNF has a bioactivity on its own (Je et al., 2012; Mizui et al., 2017; Zanin et al., 2017) whereas so far little was attempted to characterize the properties of the NGF pro-domain and how they relate to the properties of the same region within the context of proNGF (Clewes et al., 2008).

Solving the structure of proNGF has been a challenge for a long time due to the unfavourable properties of this dimeric protein. Yet, this information is very important because the pro-domain seems to play an important part in modulating or even competing with and reverting the biological effects of NGF. We approached the problem in steps. We first studied the structure of the isolated NGF pro-domain by NMR. It was previously suggested from CD studies that the isolated pro-domain has little structure in solution (Clewes et al., 2008; Kliemannel et al., 2004). We showed here that, while this is true, NGFpd contains a well-defined secondary structure propensity with regions strongly biased toward a helical conformation. We then demonstrated that the known growth cone collapse activity of proNGF (Deinhardt et al., 2011) is mainly carried out by the pro-domain, since the latter was found, surprisingly, to elicit the same response also in isolation. This activity of NGFpd appears to be p75^NTR^-dependent as it is greater in p75^NTR^ overexpressing neurons. This is an important finding which demonstrates properties specific to the pro-domain and extends the range of its reported biological activities (Clewes et al., 2008).

Building on this evidence, we approached the question of whether the isolated NGFpd is able to interact with NGF. This was not obvious: previous SAXS studies had demonstrated that the pro-domain collapses in solution into a compact structure when in the context of proNGF and significantly influences the proNGF NMR spectrum (Paoletti et al., 2009 and 2011). However, no evidence was available to demonstrate whether the interaction is persistent or transient as in a molten globule state. We were able to observe interactions between the NGFpd and NGF using a combination of NMR methods. We found that, albeit weaker than what these interactions should be in full-length proNGF, the interactions obtained by PRE studies indicate a persistent positioning of the pro-domain. Using the information obtained, we reconstructed an experimentally based model of the NGFpd-NGF adduct which could be validated both by SAXS and independent NMR data. We could thus greatly limit the conformational space explored by proNGF and trace the interactions of the pro-domain with NGF. Our MD models of proNGF suggest the presence in solution of more compact conformers in which the pro-domain interacts only with the loops of its own NGF protomer in equilibrium with more expanded conformers interacting with both NGF protomers. The latter models are consistent with and explain our previous differential scanning calorimetry measurements which showed that the proNGF dimer is thermodynamically more stable than the NGF dimer (Paoletti et al., 2011).

Our data are consistent with our previous Surface Plasmon Resonance (SPR) binding studies of proNGF towards the NGF antibodies alphaD11 and 4C8, which show a reduced affinity with proNGF as compared to NGF (Paoletti et al., 2009). They also justify a reduced affinity of proNGF to both TrkA and p75NTR receptors (Paoletti et al., 2009) and indicate a conformational heterogeneity of the pro-domains in each proNGF in contrast to the previously reported models based on HDX-MS data (Trabjerg et al., 2017).

We also obtained information on how the pro-domain influences the NGF loops in proNGF. By comparing the NMR spectra of NGF and proNGF we observed main perturbations in loops I, II and V with some minor effects on loop III and residues in the main stem of NGF induced by the pro-domain residues V87-T91. This region is the same that was demonstrated in the classic paper by Suter et al (1991) to be necessary and sufficient for the biosynthesis of correctly processed and biologically active NGF. Our data also agree with HDX-MS experiments (Trabjerg et al., 2017) which have suggested that residues R81-F89 change the structural dynamics of loops I (G144-E156), II (A161-F170) and V (D214-A219). However, in contrast with the HDX-MS experiments, our models propose the occurrence of an interaction between the exposed W142 of NGF with the pro-domain residues Y21-E29, as previously reported (Paoletti et al. 2011; Paoletti et al. 2009; Kliemannel et al. 2007). We may suggest that the V87-F91 region of the pro-domain could induce a conformational opening of loop II of the NGF domain and result in accessibility of the otherwise inaccessible crevice delimited by loops I, II and V as observed in the crystal structures of mouse NGF in the complexes with lysophosphatidylinositol (Sun and Jiang, 2015) and lysophosphatidylserine (Tong et al., 2012).

Altogether our data open a new chapter in the full understanding of the pro-domain both in isolation and within the proNGF context and suggest new directions for further investigations.

## Acknowledgments

We gratefully acknowledge Drs Geoff Kelly, Cesira De Chiara and Francesca Malerba for technical assistance and helpful discussions. AC (SNS) and AP (KCL) were supported by the collaborative EU FPT grant PAINCAGE (GA 603191) and by a Dementia Research grant. P.V.K acknowledges the support from Federal Agency of Scientific Organizations (Agreement No 007-ГЗ/Ч3363/26). AK and FLG acknowledge the support of EPSRC [grant no. EP/P022138/1; EP/P011306/1; EP/M013898/1]. HecBiosim [grant no. EP/R029407/1], PRACE and BSC’s – MareNostrum are acknowledged for computer time.

